# Comprehensive O-GlcNAc glycoproteomics on NOTCH1 EGF repeats implicated unique Lewis X epitopes in mammals

**DOI:** 10.1101/2021.11.02.463467

**Authors:** Yohei Tsukamoto, Mitsutaka Ogawa, Kentarou Yogi, Hideyuki Takeuchi, Tetsuya Okajima

## Abstract

The O-GlcNAc modification of Notch receptors regulates Notch ligand interactions in a manner distinct from other forms of O-glycans on epidermal growth factor-like (EGF) repeats of Notch receptors. Although many proteins, besides Notch receptors, are expected to be O-GlcNAcylated by EGF domain-specific O-GlcNAc transferase (EOGT), only a small number of proteins have been reported to be modified *in vivo*, and elongated O-GlcNAc glycans have not been extensively explored. To extend our view of the specificity and variety of the glycan modification, we conducted a comprehensive analysis of O-GlcNAc glycans on NOTCH1 in mammals. Mass spectrometric analysis of NOTCH1 fragments expressed in HEK293T cells revealed that several EGF domains with putative O-GlcNAcylation sites were hardly modified with O-GlcNAc. Although amino acid residues before the modification site are preferentially occupied with aromatic residues, Phe and Tyr are preferable to Trp for the apparent modification with O-GlcNAc. Furthermore, a minor form of fucosylated O-GlcNAc glycans was detected in a subset of EGF domains. Fucosylation of O-GlcNAc glycans was enhanced by *FUT1*, *FUT2*, or *FUT9* expression. The FUT9-dependent Lewis X epitope was confirmed by immunoblotting using an anti-Lewis X antibody. As expected from the similarity in the glycan structures, the Lexis X antigen was detected on O-fucose glycans. Our results refined the putative consensus sequence for the EOGT-dependent extracellular O-GlcNAc modification in mammals and revealed the structural diversity of functional Notch O-glycans.

## Introduction

O-GlcNAc modification is a unique type of post-translational modification that occurs in intracellular proteins and a limited number of extracellular proteins (Hart, G.W., Housley, M.P., et al. 2007, Ogawa, M. and Okajima, T. 2019). Extracellular O-GlcNAc occurs specifically in the epidermal growth factor-like (EGF) domain, often concomitant with O-fucose or O-glucose (Holdener, B.C. and Haltiwanger, R.S. 2019, Yu, H. and Takeuchi, H. 2019). The glycosyltransferase responsible for this modification is EOGT, the endoplasmic reticulum (ER)-localized O-GlcNAc transferase (Alam, S.M.D., Tsukamoto, Y., et al. 2020a, Matsuura, A., Ito, M., et al. 2008, Sakaidani, Y., Ichiyanagi, N., et al. 2012, Sakaidani, Y., Nomura, T., et al. 2011, Varshney, S. and Stanley, P. 2017). More than ten proteins, including Notch1 and thrombospondin-1 (TSP-1), are substrates of EOGT in mammals (Alfaro, J.F., Gong, C.X., et al. 2012, Hoffmann, B.R., Liu, Y., et al. 2012, Ogawa, M., Sawaguchi, S., et al. 2015a, Pennarubia, F., Germot, A., et al. 2021, Xu, S., Sun, F., et al. 2020). The biological importance of extracellular O-GlcNAc has been proven in studies of *Eogt*-deficient mice, which showed abnormalities in retinal angiogenesis and *EOGT* mutations in patients with Adams–Oliver syndrome (AOS), which is characterized by abnormalities in the scalp and parts of the skull (Cohen, I., Silberstein, E., et al. 2014, Ogawa, M., Sawaguchi, S., et al. 2015b, Sawaguchi, S., Varshney, S., et al. 2017, Shaheen, R., Aglan, M., et al. 2013). Furthermore, extracellular O-GlcNAc is reported to exert its function in other biological contexts, including regulatory T cell differentiation in autoimmune hepatitis (Hao, X., Li, Y., et al. 2018) and the regulation of Adropin expression in the decidualizing human endometrium (Muter, J., Alam, M.T., et al. 2018). The molecular function of O-GlcNAc has been analyzed using Notch receptors as model proteins (Ogawa, M., Tashima, Y., et al. 2020); however, many other proteins are expected to be modified by EOGT (Muller, R., Jenny, A., et al. 2013, Pennarubia, F., Germot, A., et al. 2021).

O-GlcNAcylation of EGF domains occurs on Thr/Ser residues located between the 5th and 6th conserved cysteines (Matsuura, A., Ito, M., et al. 2008). Previous research has shown that various EGF domains of NOTCH1 harboring modifiable Thr/Ser residues are O-GlcNAcylated at highly variable stoichiometry, suggesting the presence of specific sequons for preferential O-GlcNAcylation (Ogawa, M., Senoo, Y., et al. 2018). However, understanding the sequons for EOGT-dependent O-GlcNAcylation is hampered by a lack of comparative and quantitative analyses of a large number of EGF domains with putative O-GlcNAcylation sites.

In this study, we performed a comprehensive glycoproteomic analysis of O-GlcNAc glycans displayed on the entire EGF repeat of NOTCH1. The detection and semi-quantification of all the EGF domains of mouse NOTCH1 containing putative O-GlcNAcylation sites revealed that specific amino acid residues before the modification sites are critical for EOGT-dependent O-GlcNAcylation at high stoichiometry. Unexpectedly, this study led to the discovery of novel O-GlcNAc glycan glycoforms with non-reducing terminal fucose, one of which was determined to be O-linked Lewis X. Moreover, the presence of a related structure was found in O-fucose glycans. The precise mapping of O-GlcNAcylation sites and modified glycan structures on NOTCH1 provides a blueprint for the identification of other glycoproteins with extracellular O-GlcNAc and the determination of the structures of O-GlcNAc glycans on mammalian extracellular proteins.

## Results

### Comprehensive analysis of O-GlcNAcylation sites on NOTCH1 EGF repeats

Prior LC-MS/MS analysis of tryptic digests of mouse NOTCH1 EGF repeats expressed in HEK293T cells revealed only half of the 22 potential O-GlcNAcylation sites (Ogawa, M., Senoo, Y., et al. 2018). The other sites remained uncharacterized due to the longer peptide lengths of NOTCH1 trypsin digests containing potential O-GlcNAcylation sites, which often increase the structural complexity due to the co-existing O-Fuc and O-Glc glycans. However, our group and others have reported a comprehensive analysis of whole O-Glc and O-Fuc sites in the same mouse NOTCH1 expressed in HEK293T cells (Kakuda, S. and Haltiwanger, R.S. 2017, Urata, Y., Saiki, W., et al. 2020). In particular, our study showed that the 16 EGF domains (EGF2, 4, 10, 12, 13, 14, 16, 19, 20, 21, 25, 27, 28, 31, 33) with putative O-Glc sites were O-glucosylated, except for EGF9 (Urata, Y., Saiki, W., et al. 2020). The majority of O-glucosylated domains were further modified with one or two residues of Xyl, whereas EGF27 and 28 were poorly modified with Xyl. Notably, O-glucose on EGF10 is modified either with xyloses or sialyl-hexose (Urata, Y., Saiki, W., et al. 2020). Regarding O-Fuc glycans, Kakuda and Haltiwanger reported that 17 out of 20 predicted modification sites were O-fucosylated at high stoichiometry (Kakuda, S. and Haltiwanger, R.S. 2017). Among these domains, EGF6, 8, 9, 12, 26, 27, 30, 35, and 36 are modified with GlcNAc by a Fringe family of glycosyltransferases (L-FNG, M-FNG, R-FNG) and other domains (EGF2, 3, 5, 16, 18, 20, 21, 24, and 31) are poorly modified or unmodified with GlcNAc. Consistent with this report, we confirmed that EGF2, 21, and 36 were exclusively modified with O-Fuc monosaccharides (data not shown). Knowledge of the carbohydrate composition on O-glycosylation sites adjacent to the O-GlcNAcylation sites served to determine the potential O-GlcNAc glycan structures.

To cover all the O-GlcNAcylation sites on NOTCH1, NOTCH1 EGF repeats isolated from HEK293T cells were digested with either trypsin, chymotrypsin, or V8 protease. In particular, chymotrypsin is a versatile enzyme for mapping the undetermined O-GlcNAcylation sites because it cleaves the peptide bonds formed by aromatic residues that frequently occupy the −1 position from the putative EOGT modification site. The combined results from three proteases revealed that EGF domains with Tyr/Phe at the −1 position are O-GlcNAcylated at variable stoichiometry, as observed in EGF2, 10, 14, 15, 16, 20, 21, 23, 26, 27, 29, and 35, with the exception of EGF11 and 17. In contrast, all the EGF domains with Trp at the −1 position (i.e., EGF3, 7, 8, 19, and 28) were hardly O-GlcNAcylated in the wild-type HEK293T cells (**Fig. 1*A***). However, it remained possible that O-GlcNAc glycans at the chymotrypsin cleavage sites may adversely affect the cleavage efficiency of chymotrypsin and thus underestimate the O-GlcNAcylation stoichiometry. To exclude this possibility, O-GlcNAc glycans on EGF 3 and 23 analyzed through chymotrypsin digestion were independently evaluated using trypsin digestion (**Fig. 1*B***). LC-MS/MS data indicated that the O-GlcNAc glycan profiles obtained by trypsin and chymotrypsin digestion were comparable. Overall, these data suggest that EOGT favors Tyr/Phe over Trp for O-GlcNAc modification of EGF domains.

**Fig. 1.**
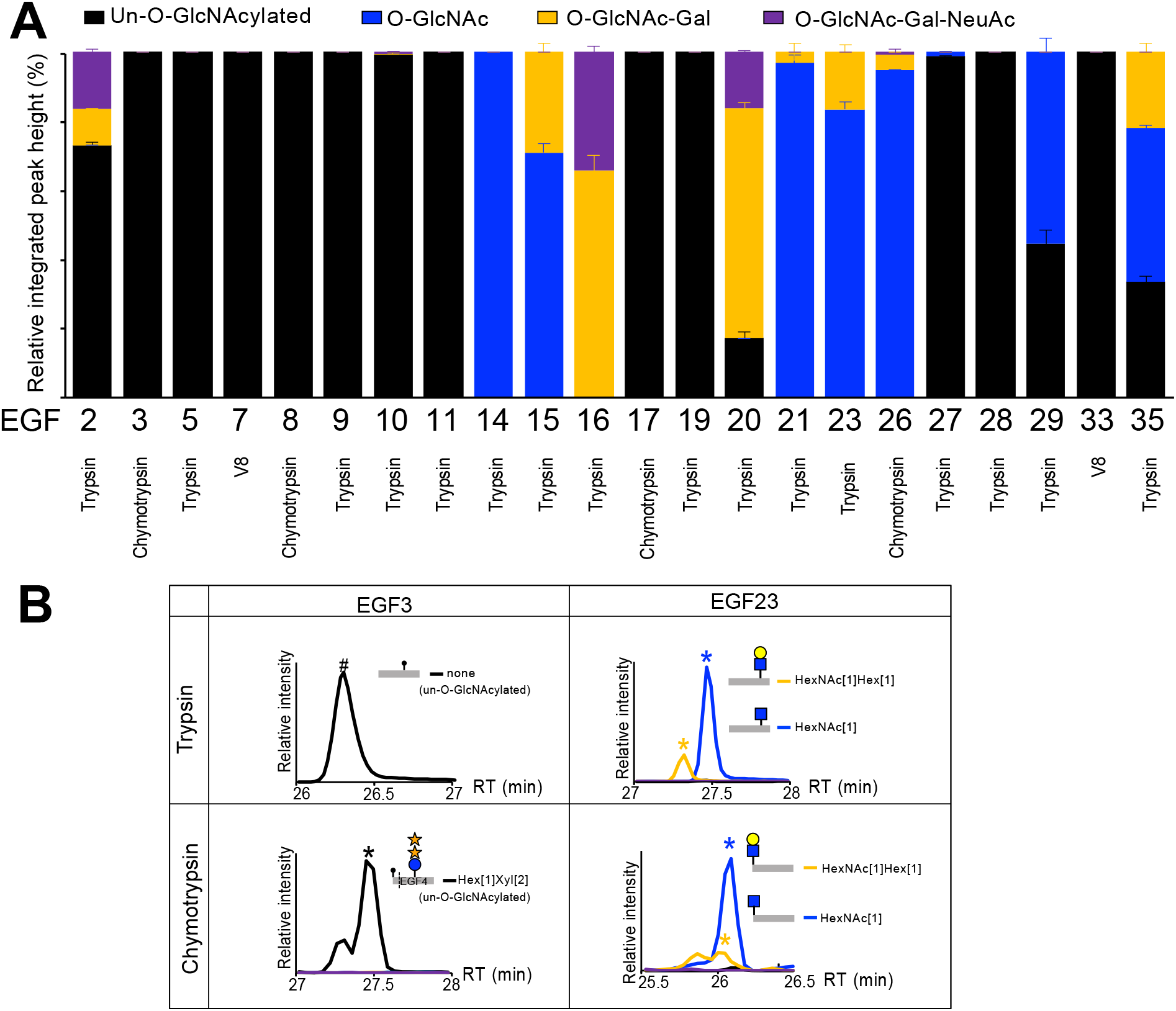
Comprehensive analysis of O-GlcNAcylation state on NOTCH1 EGF repeats. **(A)** Comprehensive glycoproteomic analysis of O-GlcNAc glycans on Notch1-EGF[1-36]:MycHis expressed in HEK293T cells. Notch1-EGF[1-36]:MycHis was digested with the indicated proteases, including trypsin, chymotrypsin, or V8, followed by LC-MS/MS analysis for semi-quantification. The stacked bar charts represent the abundance of peptides unmodified with O-GlcNAc glycans (*black*) or modified with O-GlcNAc (*blue*), O-GlcNAc-Gal (*orange*), or O-GlcNAc-Gal-NeuAc (*purple*). The proportion of integrated peak height values are presented as the mean ± the range of data (n=2). Note that semi-quantification is based on the EIC height values shown in **supplemental Table S2**. **(B)** Extracted ion chromatograms (EICs) show the relative ion intensities of protease digests containing putative O-GlcNAcylation sites at EGF3 (C^5^PPGW<u>S</u>GKSC^6^) or EGF23 (C^5^QAGY<u>T</u>GRNC^6^). Each line graph corresponds to peptides unmodified with O-GlcNAc glycans (*black*) or modified with O-GlcNAc (*blue*), O-GlcNAc-Gal (*orange*), or O-GlcNAc-Gal-NeuAc (*purple*) glycans.

The combination of proteases successfully covered all putative EOGT modification sites on NOTCH1. Consistent with previous preliminary results, only a limited number of NOTCH1 EGF domains displayed elongated O-GlcNAc-Gal-NeuAc structures (i.e., EGF2, 16, and 20). Other O-GlcNAcylated domains were not extended or poorly extended with Gal, followed by NeuAc. Importantly, this study also revealed that O-GlcNAcylation clusters in the middle part of NOTCH1 EGF repeats and the N-terminal EGF repeats are nearly devoid of O-GlcNAcylation, except for EGF2. Our results provide an overall view of O-GlcNAcylation states of all O-GlcNAcylation sites on mouse NOTCH1 under the same experimental conditions for the first time, allowing for the refinement of the requirement for O-GlcNAcylation of EGF domains.

### Identification of novel O-GlcNAc glycans containing a fucose residue

The extracellular O-GlcNAc was initially identified on *Drosophila* Notch EGF20 (dN-EGF20) produced in *Drosophila* S2 cells (Matsuura, A., Ito, M., et al. 2008). A previous study also showed that dN-EGF20 served as a substrate for mouse EOGT *in vitro* (Sakaidani, Y., Nomura, T., et al. 2011). As predicted from these results, dN-EGF20 modified with elongated structures of O-GlcNAc-Gal and O-GlcNAc-Gal-NeuAc were detected when expressed in HEK293T cells (Ogawa, M., Senoo, Y., et al. 2018). However, there have been no extensive studies investigating the structural diversity of O-GlcNAc glycans.

Using an ultrafleXtreme MALDI-TOF/TOF mass spectrometer with enhanced resolution, a mutant form of *Drosophila* Notch EGF20 lacking O-fucosylation and O-glucosylation sites but retaining O-GlcNAcylation sites (dN-EGF20^ΔGlcΔFuc^) was analyzed to monitor the glycan repertoire (**Fig. 2A**). As expected, the MALDI-TOF-MS spectra of dEGF20^ΔGlcΔFuc^ expressed in HEK293T cells indicated the presence of HexNAc[1]Hex[1] and HexNAc[1]Hex[1]NeuAc[1] (i.e., O-GlcNAc-Gal and O-GlcNAc-Gal-NeuAc). However, upon transfection with *Eogt*, a minor peak at *m/z* 8129, corresponding to HexNAc[1]Hex[1]Fuc[1], was also observed (**Fig. 2*B***). These data indicate the presence of a novel form of O-GlcNAc glycan that contains a Fuc residue.

**Fig. 2.**
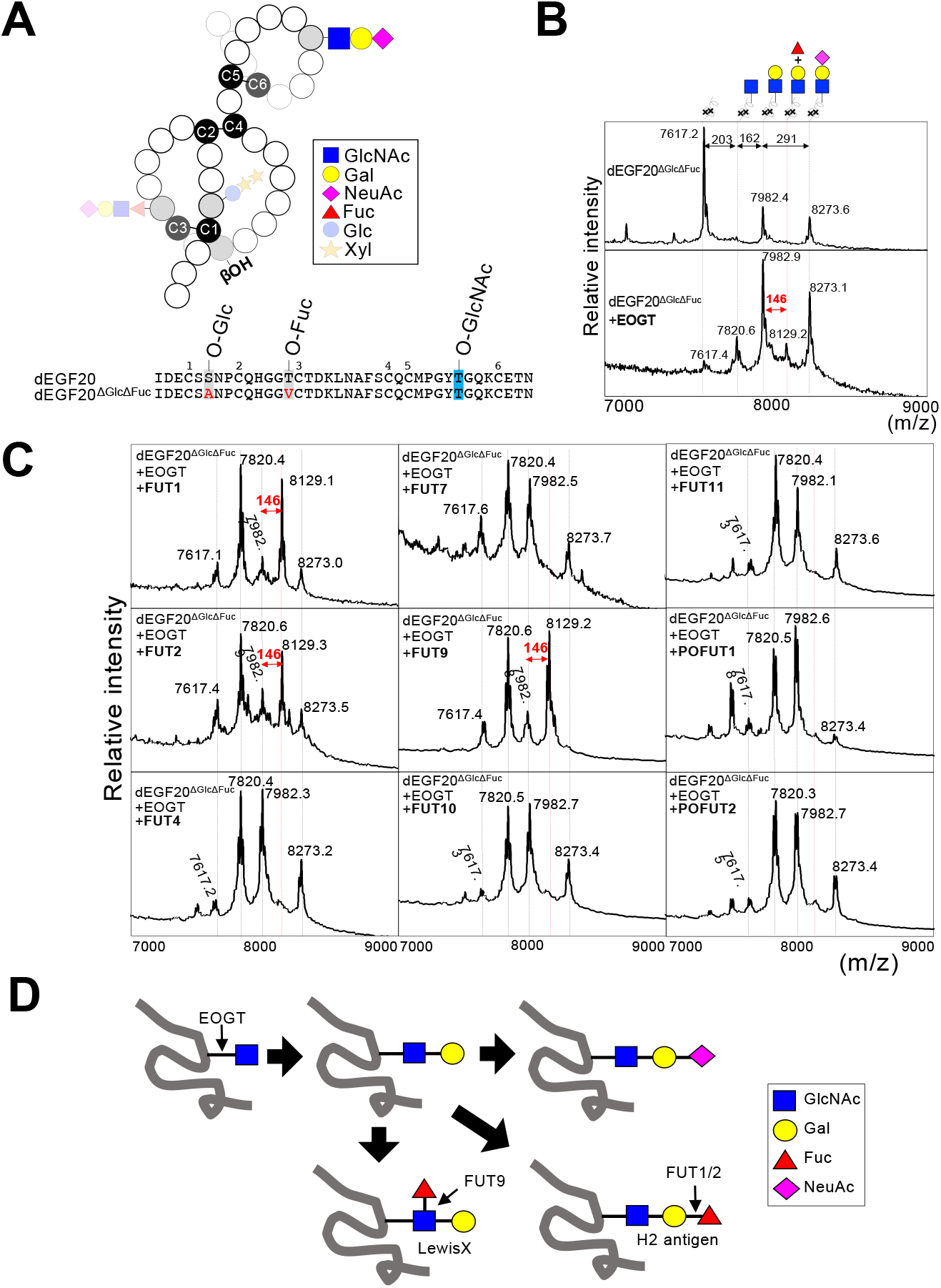
Identification of fucosyltransferases that modify O-GlcNAc glycans. **(A)** Schematic diagram of *Drosophila* Notch EGF20^ΔGlcΔFuc^ (dEGF20^ΔGlcΔFuc^) that lacks O-glucose and O-fucose sites but contains O-GlcNAc site. **(B)** dEGF20^ΔGlcΔFuc^-MycHis were transiently expressed in HEK293T cells with or without EOGT and purified using anti-His magnetic beads from the culture medium for MALDI-TOF-MS analysis. The molecular weight of the major peaks is indicated. Double-headed arrows show peak-to-peak mass increments corresponding to sugar moieties. The predicted glycans are shown above. **(C)** dEGF20^ΔGlcΔFuc^-MycHis and indicated glycosyltransferases were transiently expressed in HEK293T cells, and purified dEGF20^ΔGlcΔFuc^-MycHis was analyzed by MALDI-TOF-MS. The molecular weight of the major peaks and peak-to-peak mass increments corresponding to sugar moieties are indicated as described in (A). **(D)** The proposed structures of O-GlcNAc glycans on EGF domains and glycosyltransferases involved in their biosynthesis.

### Identification of fucosyltransferases modifying O-GlcNAc glycans

A fucosyltransferase family consists of more than 10 members of enzymes that exhibit α1,2-, α1,3-, α1,4-, α1,6-, or O-linked fucosyltransferase activity (Aplin, J.D. and Jones, C.J. 2012, Kikuchi, N. and Narimatsu, H. 2006, Schneider, M., Al-Shareffi, E., et al. 2017, Yamakawa, T., Ayukawa, T., et al. 2012). To explore the fucosylation mechanisms related to fucose-containing O-GlcNAc glycans on dN-EGF20, we searched for mouse fucosyltransferase (FUT) genes whose expression induces an increase in fucosylated O-GlcNAc glycans. As shown in **Fig. 2*C***, the expression of EOGT with FUT4, FUT7, FUT10, FUT11, POFUT1, or POFUT2 failed to induce changes in the signal at *m/z* 8129 corresponding to the HexNAc[1]Hex[1]Fuc[1] glycoform. In contrast, the co-expression of either one of the two α1,2-fucosyltransferases (FUT1 and FUT2) or α1,3-fucosyltransferase (FUT9) profoundly enhanced the *m/z* 8129 peak, concomitant with a decrease in the *m/z* 7982 peak corresponding to HexNAc[1]Hex[1] glycoform and the *m/z* 8273 peak corresponding to HexNAc[1]Hex[1]NeuAc[1]. The effect of FUT1, FUT2, and FUT9 on the expression of fucose-containing O-GlcNAc glycans was observed regardless of EOGT overexpression (**supplemental Fig. S3*A***). These data indicate that the fucosyltransferases and sialyltransferases compete for the same acceptor substrate HexNAc[1]Hex[1] and that FUT1/2 or FUT9 fucosylated HexNAc[1]Hex[1] glycan to generate H2 antigen or Lewis X antigen, respectively (**Fig. 2*D***).

In addition to Lewis X, FUT9 synthesizes Lewis Y by transferring α1,3 fucose onto the H2 antigen *in vitro* (Cailleau-Thomas, A., Coullin, P., et al. 2000). To determine whether FUT9 and FUT1/2 cooperate to generate Lewis Y structures on O-GlcNAc glycans, dN-EGF20^ΔGlcΔFuc^ was co-expressed in HEK293T cells expressing EOGT along with FUT1 and 9 or FUT2 and 9. However, the predicted HexNAc[1]Hex[1]Fuc[2] glycoform corresponding to Lewis Y was not observed (**supplemental Fig. S3*B***). Taken together, these results show that α1,2- and α1,3-fucosylation are likely to occur in a mutually exclusive manner.

### Detection of fucosylated O-GlcNAc glycans on mouse NOTCH1

Based on the results from dEGF20, we investigated the presence of fucose-containing O-GlcNAc glycans on mouse NOTCH1 EGF repeats. In addition to full-length NOTCH1, truncated forms of NOTCH1 harboring EGF1-18 were analyzed by taking advantage of the fact that deletion of C-terminal NOTCH1 increases the amount of proteins harvested from culture media, thus improving the signal-to-noise ratio in the mass spectrometry analysis.

The dataset for NOTCH1 EGF1-18 obtained in a previous study (Urata, Y., Saiki, W., et al. 2020) revealed the presence of fucosylated O-GlcNAc glycans on EGF2 and EGF16, which were unambiguously identified by the presence of oxonium ions corresponding to HexNAc (*m/z* 204), HexNAcHex (*m/z* 366), and HexNAcHexFuc (*m/z* 512) **(Fig. 3)**. It should be noted that the MS/MS spectra of glycopeptides with elongated O-Fuc glycans (i.e., O-Fuc-HexNAc-Hex) failed to produce HexNAcHexFuc oxonium ions, as stated below.

**Fig. 3.**
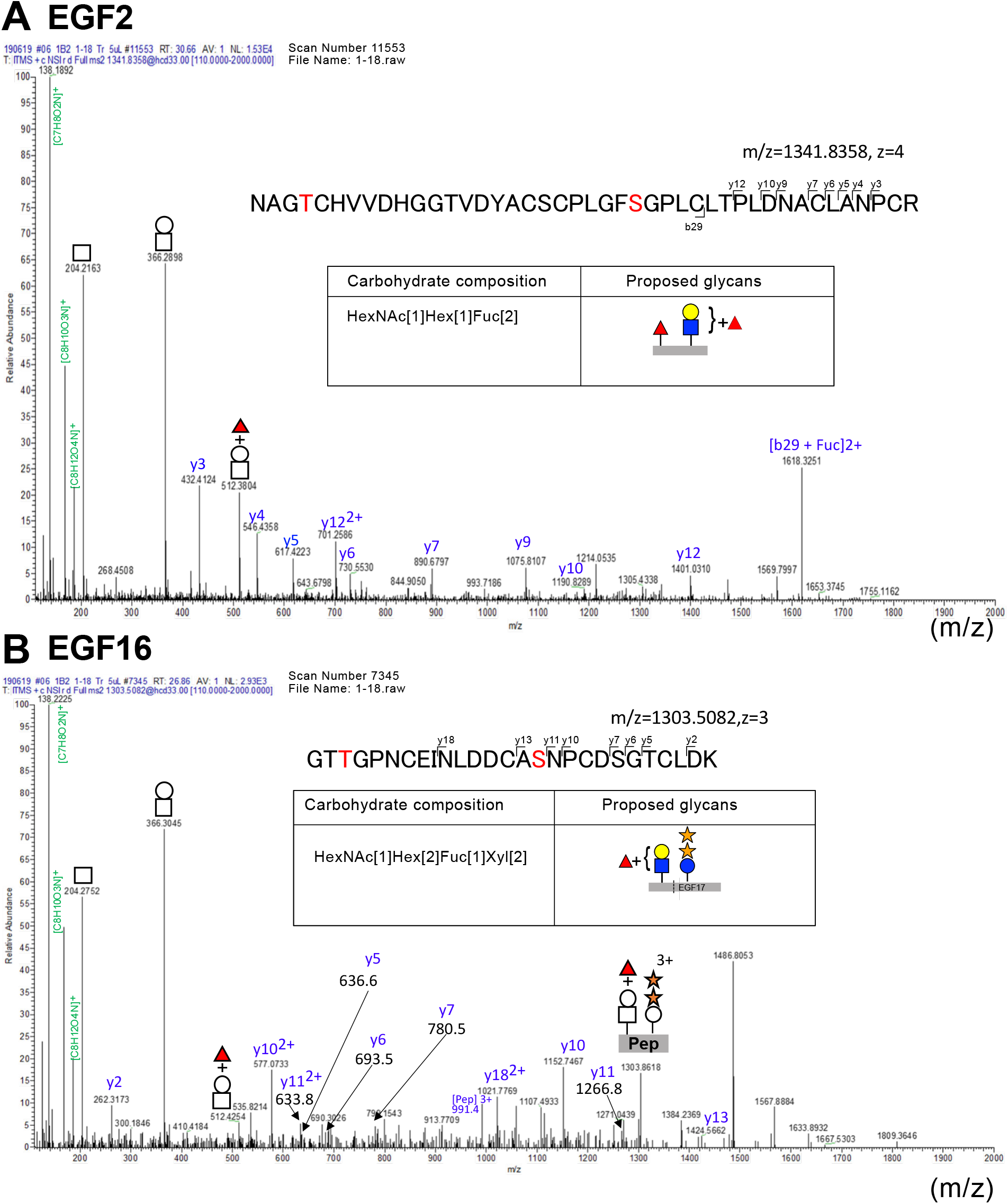
MS/MS spectra of NOTCH1 glycopeptides containing fucosylated O-GlcNAc glycans. **(A)** MS/MS spectra of glycopeptides modified with HexNAc[1]Hex[1]Fuc[1] at EGF2. NOTCH1[EGF1-18]:MycHis was expressed in HEK293T cells, and tryptic digests were analyzed by LC-MS/MS. *y* or *b* ions are indicated in blue. Proposed glycan structures were deduced based on the carbohydrate composition and knowledge of the structures of other O-glycans, as detailed in the main text. Saccharide oxonium ions are indicated in *green*. Note that the correct assignment of O-Fuc to an O-Fuc site is based on the data that EGF2 is fully occupied by O-Fuc monosaccharides. **(B)** The same with (A) but at EGF16.

Regarding fucosylated O-GlcNAc glycans on EGF2, the carbohydrate composition of total O-glycans derived from tryptic glycopeptides was HexNAc[1]Hex[1]Fuc[2] **(Fig. 3*A*)**. The glycopeptides also contain an O-Fuc site in the same EGF domain. Consistent with a previous report, chymotryptic digestion, in which the EGF2 O-Fuc site was separated from adjacent O-GlcNAc sites, confirmed the constitutive O-Fuc monosaccharide modification and thus the lack of an extended form of O-Fuc glycans on EGF2 (data not shown). Based on these observations, the carbohydrate composition at the EGF2 O-GlcNAc modification sites was determined to be HexNAc[1]Hex[1]Fuc[1].

In the case of EGF16, the carbohydrate composition of total glycans including fucosylated O-GlcNAc glycans was HexNAc[1]Hex[2]Xyl[2]Fuc[1] **(Fig. 3*B***). These tryptic glycopeptides also contain an O-Glc site on EGF17. The presence of two Xyl residues indicates O-Glc-Xyl-Xyl at the O-Glc site on EGF17, as reported previously (Urata, Y., Saiki, W., et al. 2020). It should be noted that the O-Fuc site on EGF17 does not satisfy the C^2^-X-X-X-X-S/T-C^3^ consensus sequence for O-fucosylation and is supposed to be un-O-fucosylated. Although we cannot formally exclude the possibility of occurrence of rare species carrying O-Fuc on EGF17, the detection of HexNAcHexFuc oxonium ions suggested the presence of fucosylated O-GlcNAc glycans on EGF16 with HexNAc[1]Hex[1]Fuc[1] carbohydrate composition.

### Identification of EGF Domains Modified with Fucosylated O-GlcNAc Glycans

Unlike partial NOTCH1 EGF, mass spectrometric analysis of full-length NOTCH1 EGF repeats failed to confirm fucosylated O-GlcNAc glycans, excluding EGF2 found in one of duplicate samples, presumably because of the poor yield of proteins purified from culture media. We then explored whether the presence of fucosylated O-GlcNAc glycans is manifested by the expression of FUT1, 2, or FUT9. LC-MS/MS analysis revealed that co-expression of FUT1 and EOGT produced prominent EIC peaks corresponding to the fucosylated O-GlcNAc glycans on EGF2, 16, and 20, suggesting the generation of H2 antigen (**Fig. 4 *A***).

**Fig. 4.**
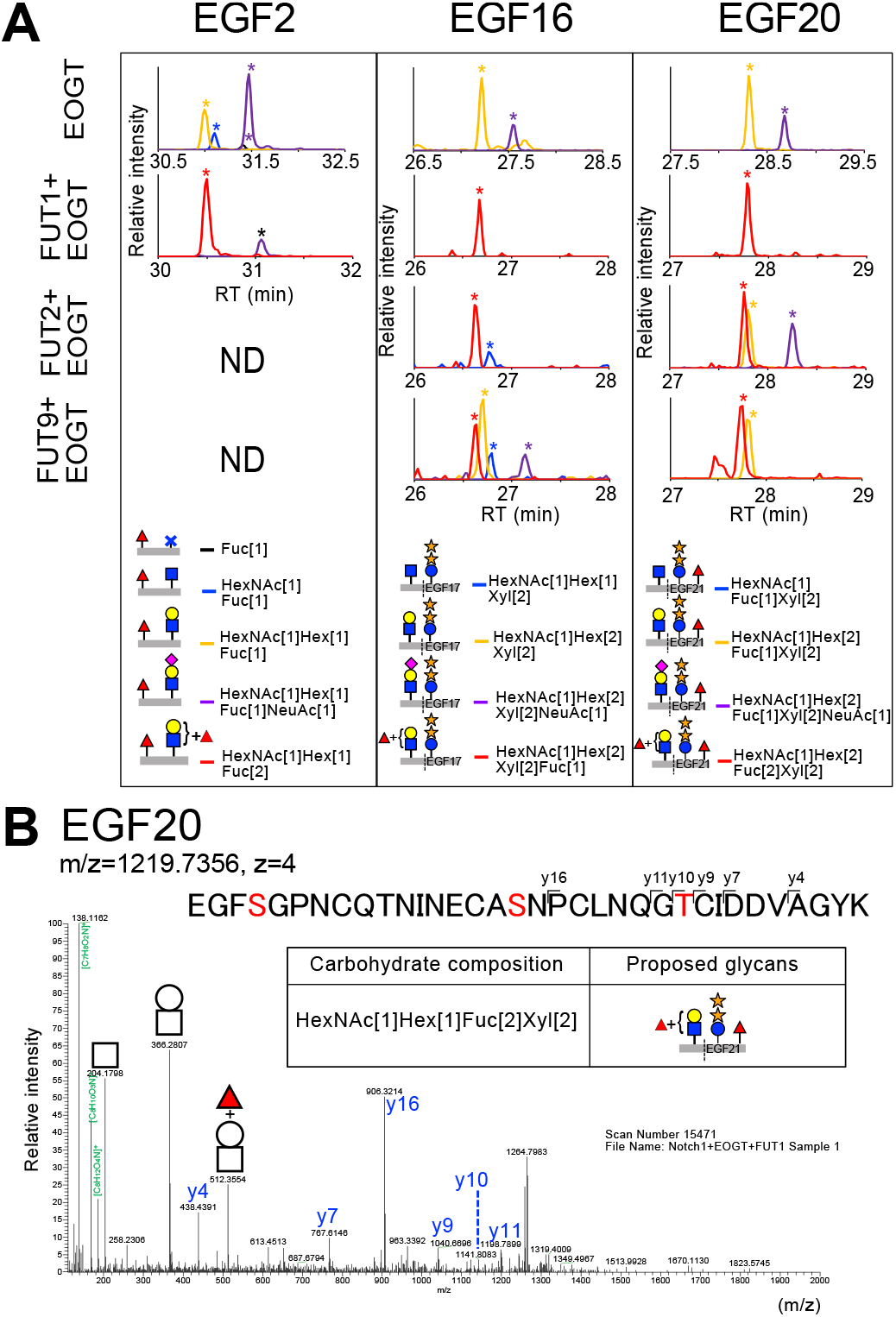
Fucosylation of O-GlcNAc glycans is enhanced by the expression of FUT1, FUT1, and FUT9. **(A)** Effect of FUT1, 2, and 9 on fucosylation of different O-GlcNAcylation sites. Full-length NOTCH1 EGF repeats were expressed in HEK293T cells along with EOGT and indicated fucosyltransferases, treated with trypsin, and analyzed by LC-MS/MS. Extracted ion chromatograms (EICs) show un-O-GlcNAcylated peptides (*black*) and glycopeptides containing O-GlcNAc (*blue*), O-GlcNAc-Gal (*orange*), O-GlcNAc-Gal-NeuAc (*purple*), or O-GlcNAc-Gal + Fuc (*red*) glycans. EIC peaks corresponding to indicated glycoforms are marked by asterisks. O-fucose is assigned based on the data that EGF2 or EGF21 is completely occupied with O-Fuc monosaccharides. Regarding data for EGF16 and 20, only the glycoforms containing O-Glc-Xyl-Xyl are displayed. **(B)** MS/MS spectra of glycopeptides modified with fucosylated O-GlcNAc with the HexNAc[1]Hex[1]Fuc[1] carbohydrate composition at EGF20. Full-length NOTCH1 EGF repeats were expressed in HEK293T cells along with EOGT and FUT1, and the tryptic digests were analyzed by LC-MS/MS. *y* or *b* ions are indicated in blue. Proposed glycan structures were deduced based on the carbohydrate composition and knowledge of the structures of other O-glycans as detailed in the main text. Saccharide oxonium ions are indicated in *green*. Note that O-fucose was manually assigned based on the data that EGF21 is fully occupied with O-Fuc monosaccharide.

As for EGF20, the carbohydrate composition of total glycans, including fucosylated O-GlcNAc, was HexNAc[1]Hex[2]Fuc[2]Xyl[2] (**Fig. 4*B***). The glycopeptides also contain O-Glc and O-Fuc sites on adjacent EGF21. The presence of two Xyl residues suggested the O-Glc-Xyl-Xyl modification at EGF21, as reported previously (Urata, Y., Saiki, W., et al. 2020). Furthermore, in agreement with a previous report (Kakuda, S. and Haltiwanger, R.S. 2017), the constitutive and exclusive O-fucosylation and the lack of the elongated forms on EGF 21 were confirmed by experiments using V8 protease, which separates O-GlcNAc and O-Fuc sites (data not shown). These observations suggest that the O-GlcNAc modification site on EGF20 carries O-glycans composed of HexNAc[1]Hex[1]Fuc[1] and that FUT1 generates H2 antigens.

As is the case of FUT1, expression of FUT2 or FUT9 confers fucosylated O-GlcNAc glycans on NOTCH1 EGF16 and 20, suggesting that these enzymes generate H2 or Lewis X antigen, respectively (**Fig. 4*A***). Taken together, these data suggest that *FUT1*, *FUT2*, and *FUT9* modify O-GlcNAc glycans in the selective EGF domains of NOTCH1.

### Identification of fucosylated O-Fuc glycans on NOTCH1 EGF26

O-Fuc monosaccharides were modified with GlcNAc by FRINGE GlcNAc transferases and β4-galactosyltransferase, leading to the formation of extended O-fucose-β1,3GlcNAc-β1,4Gal trisaccharide (Brückner, K., Perez, L., et al. 2000, Moloney, D.J., Panin, V.M., et al. 2000). The similarity between O-GlcNAc and O-fucose glycans in the shared GlcNAcβ1,4-Gal structure led us to hypothesize that O-Fuc glycans could be fucosylated by a similar mechanism. To address this possibility, the NOTCH1 EGF24-28 fragment, which includes O-fucosylated EGF domains modifiable by L-Fringe, was employed (**Fig. 5*A***). Consistent with previous reports, modifications to the EGF26 O-Fuc site were found in the form of a monosaccharide (**Fig. 5*B***) (Kakuda, S. and Haltiwanger, R.S. 2017). By expressing L-Fringe, elongated forms of O-fucose glycans were increased (**Fig. 5*C***).

**Fig. 5.**
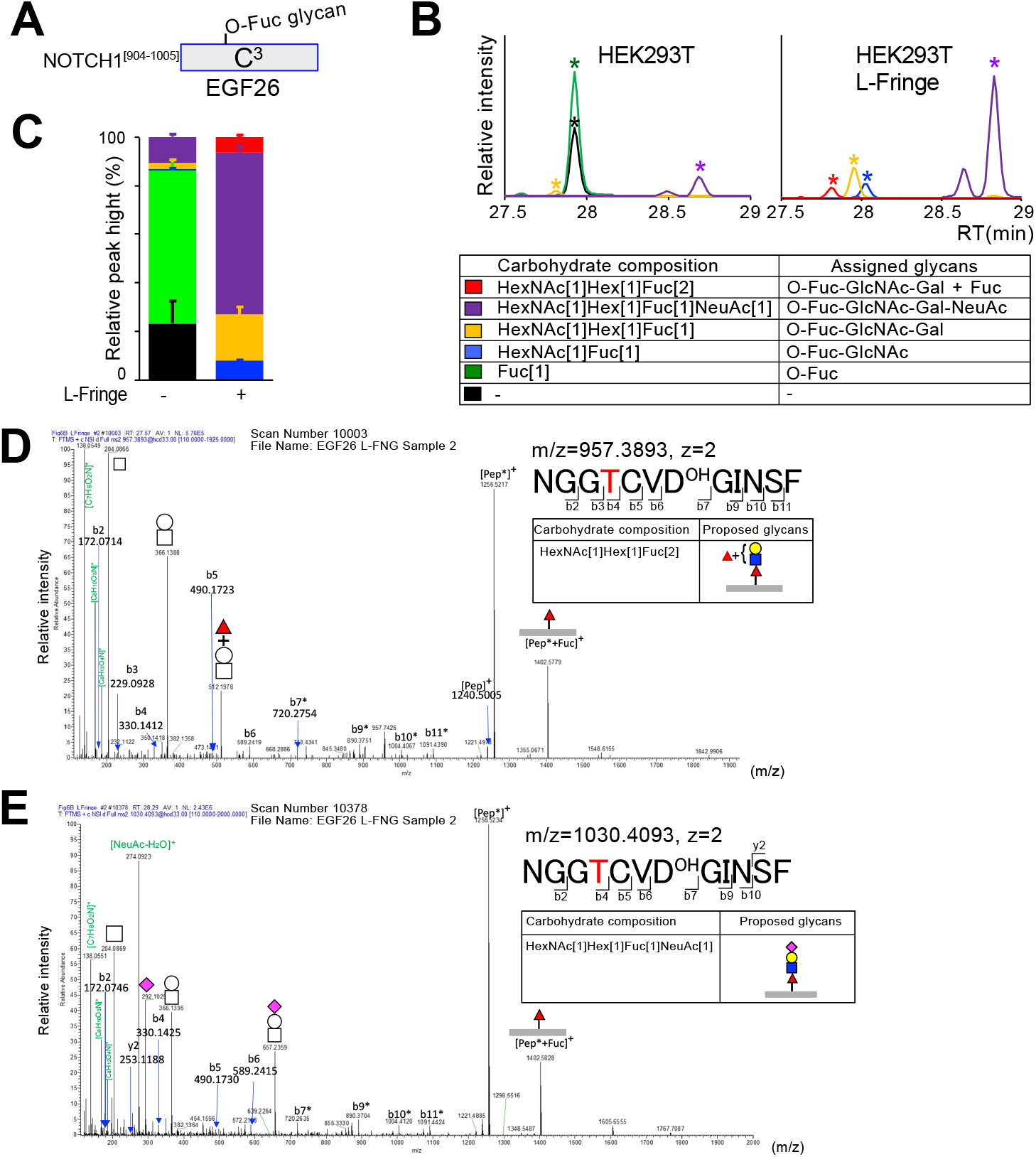
O-Fuc glycans on NOTCH1 EGF26 contain non-reducing terminal fucose. **(A)** Schematic representation of chymotryptic peptides corresponding to NOTCH1^[904-1005]^ fragment that harbors C^2^-X-X-X-X-S/T-C^3^ O-Fuc consensus sequence. **(B)** Extracted ion chromatograms (EICs) of NOTCH1^[904-1005]^ fragment carrying O-Fuc glycans. HEK293T cells were transfected to express NOTCH1-EGF[24-28] with or without L-Fringe. The chymotryptic digests of purified NOTCH1 fragments were analyzed by LC-MS/MS. Shown are relative ion intensities of NOTCH1^[904-1005]^ fragment lacking O-Fuc glycans (*black*) and glycopeptides containing O-Fuc (*green*), O-Fuc-GlcNAc (*blue*), O-Fuc-GlcNAc-Gal (*orange*), O-Fuc-GlcNAc-Gal-NeuAc (*purple*), or O-Fuc-GlcNAc-Gal + Fuc (*red*). EIC peaks corresponding to indicated glycoforms are marked by asterisks. **(C)** Semi-quantitative measurement of unglycosylated or O-fucosylated peptides derived from NOTCH1 EGF26 domain. Data are presented as the mean ± the range of the data (n=2). **(D)** MS/MS spectra of glycopeptides modified with HexNAc[1]Hex[1]Fuc[2] at EGF26. NOTCH1-EGF[24-28] was co-expressed with L-Fringe in HEK293T cells, affinity-purified, and digested with chymotrypsin for LC-MS/MS analysis. Saccharide oxonium ions are indicated in *green*. The fragment ions containing hydroxylated aspartic acid are indicated by asterisks. **(E)** MS/MS spectra of glycopeptides modified with HexNAc[1]Hex[1]Fuc[1]NeuAc[1] at EGF26. Data were obtained as described in (D).

In addition to previously identified glycoforms, O-Fuc glycans with a carbohydrate composition of HexNAc[1]Hex[1]Fuc[2] were identified (**Figure 5*B* and *C*)**. As with fucosylated O-GlcNAc glycans, oxonium ions corresponding to HexNAc (*m/z* 204), HexNAcHex (*m/z* 366), and HexNAcHexFuc (*m/z* 512) were detected (**Fig. 5*D***). In contrast, well-characterized glycoforms corresponding to O-Fuc-GlcNAc-Gal and O-Fuc-GlcNAc-Gal-NeuAc failed to produce HexNAcHexFuc oxonium ions (**Fig. 5*E*** and **supplemental Fig. S3**). These results indicate that HexNAcHexFuc oxonium serves as a diagnostic ion for novel fucosylated glycans at O-Fuc and O-GlcNAc sites. Notably, O-fucosylated glycopeptides carrying sialyl Lewis X epitopes (HexNAc[1]Hex[1]Fuc[2]NeuAc[1]) or Lewis Y epitope (HexNAc[1]Hex[1]Fuc[3]) were not detected on EGF26.

### FUT9 Generates O-Lewis X Epitopes on O-GlcNAc and O-fucose Glycans

To detect the non-reducing terminal fucose on O-GlcNAc or O-Fuc glycans, NOTCH1 EGF repeats were subjected to immunoblotting. We focused on the expression of the Lewis X epitope because anti-type H antibodies suitable for the analysis were unavailable. NOTCH1 EGF repeats were expressed in wild-type HEK293T or *EOGT* mutant cells in the presence or absence of FUT9 and L-Fringe (**Fig. 6**). Only faint immunoreactivity toward the anti-Lewis X antibody was observed for NOTCH1 EGF repeats in the absence of exogenous expression of FUT9 and L-Fringe. A similar result was obtained when L-Fringe was co-expressed in these cells.

**Fig. 6.**
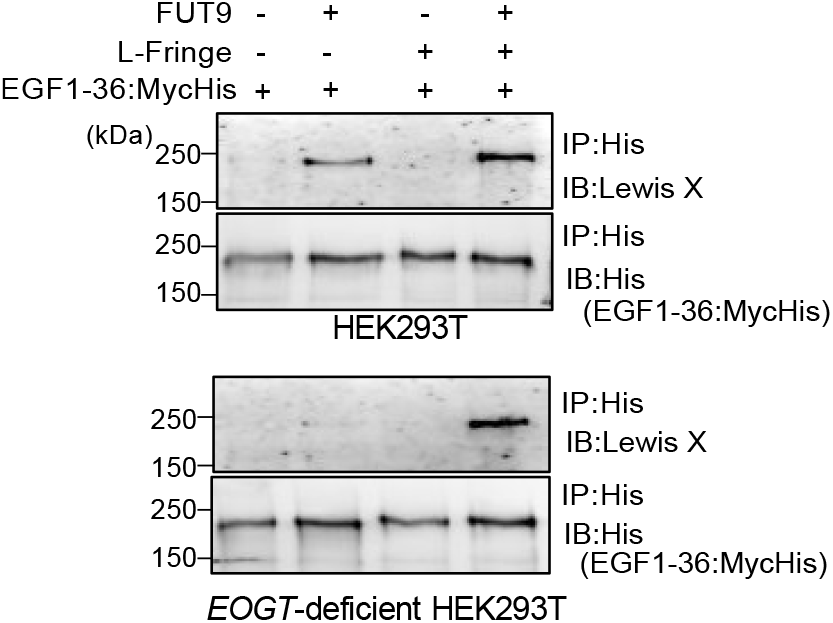
FUT9 generates Lewis X epitopes on O-GlcNAc and O-fucose glycans. NOTCH1-EGF[1-36]: MycHis was transiently expressed together with L-Fringe, FUT9, or both in HEK293T cells or *EOGT*-deficient HEK293T cells, affinity-purified with anti-His magnetic beads, and analyzed by western blotting with the indicated antibodies.

In contrast, co-expressing FUT9 conferred an elevated Lewis X signal to NOTCH1 EGF repeats in HEK293T cells but not in *EOGT* mutant cells, indicating the presence of Lewis X epitopes on O-GlcNAc glycans. In *EOGT* mutant cells, Lewis X reactivity was observed only when FUT9 was simultaneously expressed with L-Fringe, suggesting that Lewis X epitopes can be created on O-fucose glycans. Moreover, the FUT9-dependent Lewis X signal was elevated in an additive manner by co-expressing L-Fringe in wild-type HEK293T cells. These data suggest that Lewis X epitopes are present on both O-GlcNAc and O-fucose glycans on NOTCH1.

## Discussion

This study used mass spectrometry to conduct semi-quantitative measurements of O-GlcNAc glycans on all potential O-GlcNAcylation sites on NOTCH1 (**Fig. 7*A***). Previous studies reported half of the potential O-GlcNAcylation sites and basic O-GlcNAc glycan structures on NOTCH1, but the overall O-glycosylation pattern on NOTCH1 was unknown (Ogawa, M., Senoo, Y., et al. 2018). The comprehensive analysis of all the O-GlcNAcylation sites conducted in the present study showed that O-GlcNAcylation and other O-glycosylations occurred independently in the individual EGF domains (**Fig. 7*B***). Although the −1 position from the putative O-GlcNAcylation site is preferentially occupied by aromatic residues, only Phe and Tyr, not Trp, are pivotal for O-GlcNAcylation at a higher stoichiometry (**Fig. 7*A***). As reported previously, O-GlcNAcylation was barely detected on NOTCH1 EGF28, which has a Trp-Thr sequence, unless EOGT is exogenously expressed (Ogawa, M., Senoo, Y., et al. 2018). The findings presented in this study provide further evidence that other EGF domains containing Trp-Thr sequences (i.e., EGF3, 7, 8, 19) were exclusively devoid of O-GlcNAc modification in HEK293T cells. It is worth mentioning that other factors are likely to further impose the O-GlcNAcylation states of EGF domains since EGF 11 and 27 contain the Tyr-Thr consensus but are poorly O-GlcNAcylated (**Fig. 7*B***). The strict requirement for preferential O-GlcNAcylation explain why few proteins were identified as O-GlcNAcylated EGF domains in previous glycoproteomic studies in mouse brains (Alfaro, J.F., Gong, C.X., et al. 2012). The refined putative consensus sequences for preferential O-GlcNAcylation, defined as C^5^-X-X-G/S/P-Y/F/T-T/S-G-X-X-C^6^, predict approximately 100 proteins that are modified with O-GlcNAc at high stoichiometry by EOGT in mice and humans (**supplemental Fig. S4**). The preference of Phe and Tyr over Trp may be applicable to *Drosophila* NOTCH, whose EGF28 domain with C^5^XXGWTXXC^6^ sequence is devoid of O-GlcNAc modification (Harvey, B.M., Rana, N.A., et al. 2016).

**Fig. 7.**
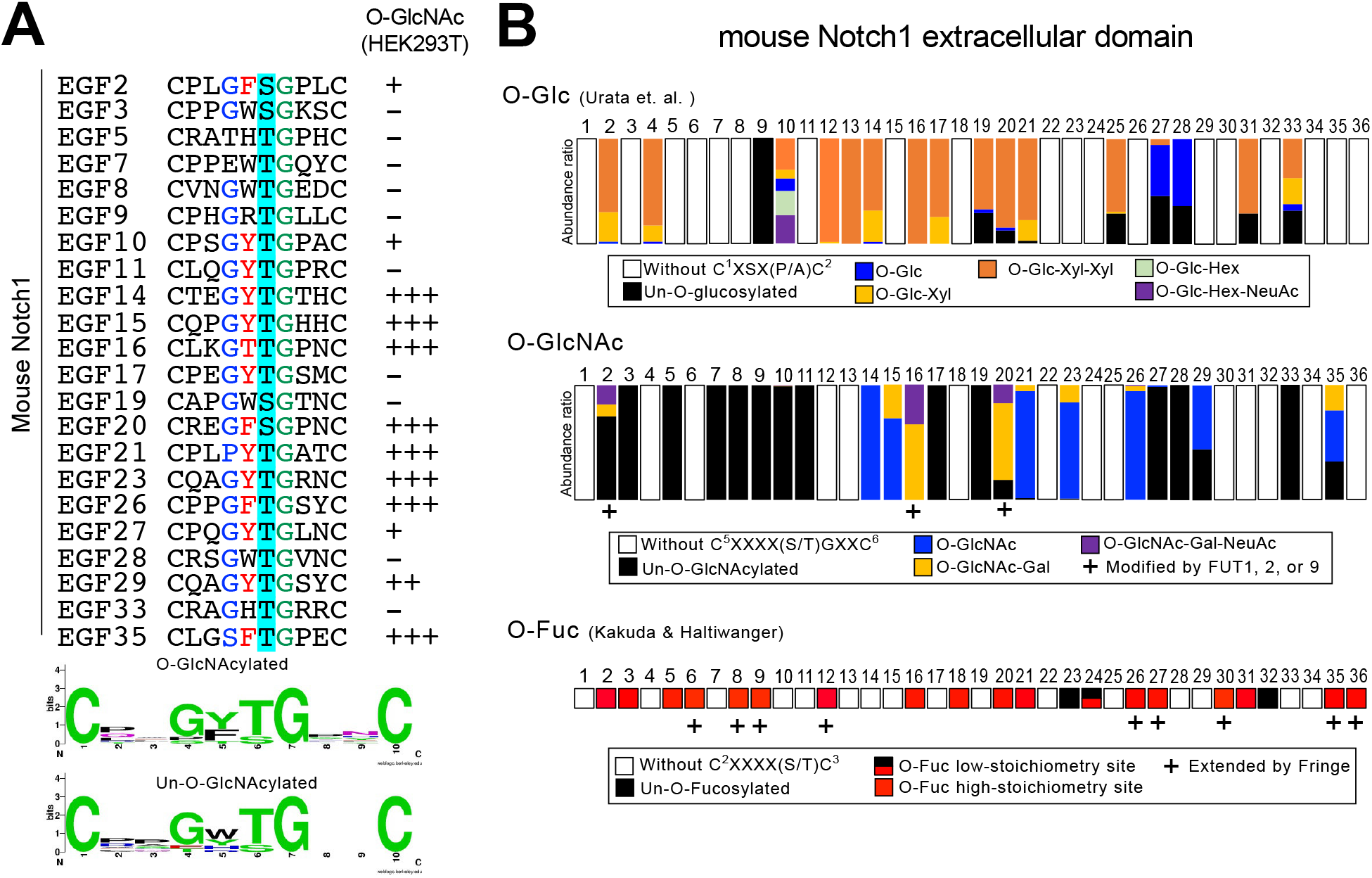
Refined consensus sequence for extracellular O-GlcNAc modification in mammals. **(A)** List of NOTCH1 EGF domains with Thr/Ser residues at the O-GlcNAcylation sites. The amino acid sequences between 5th and 6th conserved Cys residues of EGF repeats are shown. In HEK293T cells, O-GlcNAcylation occurs on EGF2, 10, 14, 15, 16, 20, 21, 23, 26, 27, 29, and 35. Note that O-GlcNAcylation was detected on NOTCH1 EGF11 and 28 when EOGT was exogenously expressed in HEK293T cells (Ogawa, M., Senoo, Y., et al. 2018). Sequence logos for O-GlcNAcylated and un-O-GlcNAcylated sequence motifs were generated by WEBLOGO3. **(B)** Schematic representation and O-glycosylation patterns on mouse NOTCH1 EGF repeats expressed in HEK293T cells. The abundance ratio for O-glycans was adopted from (Urata, Y., Saiki, W., et al. 2020) and O-fucose glycans were referred to (Kakuda, S. and Haltiwanger, R.S. 2017).

Despite the minor glycoforms, fucosylated O-GlcNAc glycans were detected on EGF2 and 16. Exogenous expression of FUT1/2 or FUT9 increased the population of the HexNAc[1]Hex[1]Fuc[1] O-GlcNAc glycoforms, concomitant with a decrease in the HexNAc[1]Hex[1] O-GlcNAc glycoform, indicating the generation of H2 antigen or Lewis X antigen, respectively (**Fig. 4*A***). NOTCH O-glycans with non-reducing terminal fucose were readily detectable on O-Fuc glycans in Fringe-expressing cells (**Fig. 5**). In Notch receptors, fucose residues are found at the reducing terminus (i.e., O-fucose), in contrast to other types of glycans that display fucose moieties at the non-reducing terminal end. However, this study revealed the modification of fucose at both the reducing and non-reducing termini of NOTCH1 O-glycans.

α1,3-fucosyltransferases are involved in synthesizing various glycan structures, and their syntheses require distinct sets of α1,3-fucosyltransferases that exhibit different substrate specificities. In humans, six α1,3-fucosyltransferases fucosylate lactosamine units to produce Lewis X or sialyl Lewis X. Among these, FUT9 and FUT4 are the most potent in the fucosylation of an unsialylated LacNAc to create Lewis X (Mondal, N., Dykstra, B., et al. 2018). All α1,3-fucosyltransferases, except FUT9, can create sialyl Lewis X (Kudo, T., Ikehara, Y., et al. 1998, Mondal, N., Dykstra, B., et al. 2018). Thus, the detection of Lewis X and sialyl LacNAc, but not sialyl Lewis X, on O-GlcNAc glycans indicates that FUT9 activity is responsible for the biosynthesis of the Lewis X epitope (**Fig. 2*D***). Accordingly, the expression of mouse FUT9, but not FUT4 and FUT7, resulted in fucosylation of O-GlcNAc glycans on dEGF20 (**Fig 2*C***). These findings indicate that the terminal fucosylation of NOTCH1 O-GlcNAc glycans is regulated by mechanisms dependent on specific α1,3-fucosyltransferases, rather than collateral processes mediated by nonspecific α1,3-fucosyltransferases.

It has been reported that FUT1 prefers GlcNAc-β1,4Gal over GlcNAc-β1,3Gal, in contrast to FUT2, which shows higher activity toward GlcNAc-β1,3Gal than GlcNAc-β1,4Gal *in vitro* (Oriol, R. and Mollicone, R. 2014). Consistent with the difference in the substrate specificities, the expression of FUT1 resulted in a marked increase in the HexNAc[1]Hex[1]Fuc[1] O-GlcNAc glycans on NOTCH1 EGF20 as compared with FUT2. FUT1 and FUT2 similarly decreased the population of HexNAc[1]Hex[1] O-GlcNAc glycans (**Fig. 4*C***), suggesting that the H2 antigen (O-GlcNAc-β1,4Galα1,2Fuc) was generated from O-LacNAc (**Fig. 2*D***). Whether endogenous α1,2-fucosyltransferase activity, mediated by FUT1/2, contributes to the expression of the H2 antigen on NOTCH1 remains to be elucidated.

The expression of FUT9 in HEK293T cells creates Lewis X epitopes on NOTCH1, depending on the presence of O-GlcNAc or O-fucose. These results indicate that α1,3-fucosyltransferase, including FUT9, simultaneously regulates the structure and function of both O-fucose and O-GlcNAc. Interestingly, the knockdown of FUT9 in neural stem cells inhibits cell proliferation and reduces the mRNA expression of *Hes5*, a Notch target gene (Yagi, H., Saito, T., et al. 2012).

In summary, this study is the first to elucidate the landscape of O-GlcNAc glycans on 36 EGF repeats of NOTCH1. A repertoire of O-GlcNAc glycan structures will serve as a blueprint for identifying structurally related EGF repeat-containing glycoproteins, which are physiologically crucial for cellular signaling and extracellular matrix function.

## Materials and methods

### Plasmid constructs

Expression vectors for NOTCH1 EGF repeats (pSectag2C/mNotch1-EGF1-36:MycHis-IRES-EGFP), EOGT (pSectag2C/EOGT), and dEGF20^ΔGlcΔFuc^ (pSectag2C/dEGF20^ΔGlcΔFuc^:MycHis-IRES-EGFP) have been previously reported (Ogawa, M., Sawaguchi, S., et al. 2015b, Ogawa, M., Senoo, Y., et al. 2018). Expression vectors for NOTCH1 EGF24-28 repeats (pSectag2C/mNotch1-EGF[24-28]:MycHis) have been reported previously (Shao, L., Moloney, D.J., et al. 2003). The expression vectors for mouse fucosyltransferases (FUTs) and L-Fringe were obtained from Origene (**supplemental Fig. S1**).

### Cell culture and stable cell line

HEK293T cells and *EOGT*-deficient HEK293T cells (Alam, S.M.D., Tsukamoto, Y., et al. 2020b) were cultured in Dulbecco’s modified Eagle’s medium (DMEM) supplemented with 10% fetal bovine serum (FBS) (Invitrogen). To express full-length or partial NOTCH1 EGF repeats, the expression vectors were transfected into HEK293T cells with or without those encoding the indicated glycosyltransferases using PEI-MAX, as described previously (Ogawa, M., Senoo, Y., et al. 2018).

### MALDI-TOF-MS analysis

Mass spectrometric analyses of dN-EGF20^ΔGlcΔFuc^-MycHis were performed using an ultrafleXtreme (Bruker). Briefly, expression vectors encoding Notch fragments were transfected into HEK293T cells using PEI-MAX in OPTI-MEM (Thermo Fisher Scientific). On the following day, the culture medium was changed to DMEM with 10% FBS. After 48 h, each culture medium was collected and centrifuged at 500 × *g* for 15 min. The resulting supernatants were incubated overnight with anti-His-tag mAb-magnetic beads (MBL). The magnetic beads were washed with PBS and eluted with 0.1% TFA in 10% acetonitrile. The samples were desalted using a Ziptip (Millipore) and dried in a vacuum freeze dryer. The dried samples were resuspended in 3 μL of 0.1% TFA in 10% acetonitrile and spotted on a MALDI plate. After drying, 1 μl of DHB (Wako) in 0.1% TFA and 10% acetonitrile was spotted onto a MALDI plate and air-dried. The MS spectra were recorded as the average of 6000–10000 laser shots, with a laser intensity ranging from 80 to 100 in linear positive ion mode.

### Hybrid quadrupole FT linear ion trap MS analysis

To purify full-length or partial NOTCH1 EGF repeats, each culture medium was incubated with anti-His-tag mAb-magnetic beads (MBL) (Ogawa, M., Senoo, Y., et al. 2018). After washing with PBS, the proteins bound onto the magnetic beads were eluted using SDS sample buffer containing 2% SDS and 70 mM 2-mercaptoethanol. For in-gel digestion, the samples were subjected to SDS-PAGE and stained with a gel-negative stain kit (Nacalai). The stained bands containing the NOTCH1 fragment were excised and transferred to low-binding protein tubes (Ina Optica). Acetonitrile was added to the gels and incubated for 10 min, followed by speed-vacuuming for 20 min for drying. To reduce and alkylate the samples, 10 mM DTT was added to the gels in 100 mM NH_4_HCO_3_ for 30 min, followed by 50 mM iodoacetamide in 100 mM NH_4_HCO_3_ for 30 min. Next, the gels were incubated with 50% MeOH in 5% acetic acid at 4°C. On the following day, acetonitrile was added to the gels, which were subsequently dried using a speed-vacuum system. The samples were digested with 1 μg/μl trypsin gold (Promega), chymotrypsin (Promega), or V8 protease (New England Biolabs) in 25 mM NH_4_HCO_3_ overnight at 37 °C. Each solution containing the digested peptides was desalted with Ziptip (Millipore) and dissolved in 10% acetonitrile in 0.1% TFA.

The peptides were analyzed using an Orbitrap Fusion Tribrid mass spectrometer (Thermo Fisher Scientific) coupled to an UltiMate3000 RSLC nano-LC system (Dionex Co.) with a nano-HPLC capillary column (150 mm × 75 μm, Nikkyo Technos) via a nanoelectrospray ion source. The LC gradient with 0.1% formic acid as solution A and 0.1% formic acid 90% acetonitrile as solution B was set as follows: 5%–100% B (0–45 min), 100%–5% B (45–45.1 min), and 5% B (45.1–60 min). Data-dependent tandem MS analysis was performed using a top-speed approach (cycle time of 3 s). Precursor ions were analyzed in an Orbitrap mass analyzer, whereas fragment ions generated by HCD fragmentation were analyzed in an Orbitrap or a linear ion trap mass analyzer. Data acquisition parameters were summarized in **supplemental Table S1**.

### Qualitative and semi-quantitative analysis of O-GlcNAc Glycan

Peak lists were generated using extract_msn in Xcalibur 3.0.6 (Thermo Scientific) with the default parameters. Data analysis was performed by using GlycoPAT software (version 1.0) (Liu, G., Cheng, K., et al. 2017). To generate *in silico* glycopeptide digest library, the following input files and parameters were specified: mouse NOTCH1 (Uniprot ID: Q01705 [Last modified April 10, 2018]) amino acid sequence [20-1426] as a peptide sequence; trypsin (specific for cleavage at the C-terminus of Lys/Arg), chymotrypsin (specific for cleavage at the C-terminus of Trp/Tyr/Phe), or V8 protease (specific for cleavage at the C-terminus of Glu/Asp) as proteolytic enzymes with the possibility of zero missed cleavage, with the exception of two missed cleavages for a chymotryptic fragment at EGF26; carbamidomethylation on Cys residues as a fixed modification; methionine oxidation as a variable modification; β-hydroxylation of Asn/Asp at EGF 7/9/11/12/13/14/15/16/17/18/19/20/21/23/25/26/27/30/31/32 that conforms to the C^3^X(<u>D/N</u>)XXXX(F/Y)XC^4^XC^5^ consensus sequence as a variable modification (Wouters, M.A., Rigoutsos, I., et al. 2005). The following glycan lists were also specified as variable modifications: {n}, {n{h}}, {n{h{s}}}, {n{h{f}}}, {n{h{s}{f}}}, {n{h{f}{f}}} at the O-GlcNAcylation sites; {h}, {h{x}}, {h{x{x}}} at the O-Glc sites except for EGF10; {h}, {h{h}}, {h{h{s}}} {h{x}}, {h{x{x}}} at the O-Glc sites on EGF10; {h} at the atypical O-Glc sites on EGF11, {f} at O-Fuc sites on EGF2, 3, 5, 16, 18, 20, 21, 31; {f}, {f{n}}, {f{n{h}}}, {f{n{h{s}}}}, {f{n{h{f}}}}, {f{n{h{f}{f}}}} {f{n{h{s}{f}}}} at O-Fuc sites on EGF6, 8, 9, 12, 26, 27, 30, 35, 36, in which Hex, HexNAc, NeuAc, Fuc, and Xyl are annotated by “h,” “n,” “s”, “f,” and “x”, respectively. MS/MS analysis and scoring was performed using the glycopeptide digest library and the following parameters: MS1 tolerance, 20 ppm; MS/MS tolerance, 0.5 Da; HCD mode. In order to view the results for manual inspection, we applied an Ensemble Score > 0.2, which corresponded to MS/MS spectra with multiple fragment ions for a reliable peptide identification. For the inspection of MS/MS spectra, correct assignment of major b/y ions and characteristic glycan-derived fragment ions at 138, 168, 186, 204, 274, 292, 366, 512, and 657 *m/z* were manually inspected using the GlycoPAT and Xcalibur Qual browser (version 4.2, Thermo Fisher Scientific). List of O-GlcNAcylated and un-O-GlcNAcylated peptides and their annotated MS/MS spectra generated by the GlycoPAT software are provided in **supplemental Tables S2 and S3**. Where specified, the proposed glycan structures were determined by integrating the information of glycoforms obtained from multiple glycopeptides generated by different proteases.

If particular glycoforms were detected only in one of the two replicates, the other replicate was subject to manual data analysis based on the expected retention time to check for reproducibility. In rare cases, β-hydroxylated or un-O-GlcNAcylated peptides were undetected by the GlycoPAT software. Manual inspection of the MS/MS spectra identified un-O-GlcNAcylated peptides at EGF7, EGF17, EGF29, and EGF33 (**supplemental Tables S4)**, and an O-GlcNAcylated/β-hydroxylated glycopeptide at EGF14 (**supplemental Tables S3)**.

The extracted ion chromatograms (EICs) were generated using the Xcalibur Qual browser by selecting the most abundant isotopic peaks (monoisotopic peak or second isotope peak) with Gaussian smoothing at five points. The EIC peak height was measured and compensated for by the isotope abundance ratio. The relative intensity derived from all the observed charge states of each glycopeptide was summed. The proportion of integrated peak height values for specified glycoform to those corresponding to all detectable (glyco-) peptides was calculated for semi-quantification.

### Western blotting of Notch1 EGF1-36:MycHis

The culture media were incubated with anti-His-tag mAb-magnetic beads (MBL). The magnetic beads were washed with PBS and eluted with SDS sample buffer containing 2% SDS and 70 mM 2-mercaptoethanol. Each sample was separated using SDS–PAGE and transferred to a polyvinylidene fluoride membrane (Millipore). Immunodetection was performed using an appropriate primary antibody, followed by an HRP-conjugated secondary antibody. The bands were visualized with Immobilon Western Chemiluminescent HRP Substrate (Millipore) and exposed to X-ray film (Fujifilm) or iBright FL1500 (Thermo). The antibodies used were anti-Lewis X antibody (1:1000) (SH1; Wako), anti-O-GlcNAc antibody (1:1000) (CTD110.6; Thermo), and anti-His antibody (1:2000) (OGHis; MBL).

### Experimental Design and Statistical Rationale

For the semi-quantification of O-GlcNAc glycans using LC-MS/MS, two biological replicate experiments were performed. Data are expressed as the mean ± the range of the data. The data obtained by MO and KY were independently confirmed by YT and TO. Unless otherwise noted, only glycopeptides detected in both of two replicates by the GlycoPAT software or manual inspection were considered to be identified and subjected to semi-quantitative analysis. Three biological replicate experiments were conducted for MALDI-MS for each sample, and representative mass spectra were presented. For immunoblotting, three biological replicate experiments were performed for each experiment, and representative images were shown.

## Acknowledgments

We thank K. Taki (Nagoya UniversityUniv) for conducting the LC-MS/MS analysis, K. Oyama (Nagoya University) for conducting the MALDI-TOF-MS analysis, K. Furukawa (Chubu University) for supervising and supporting the project, and S. Takeda (Nanzan Kokusai High School) for conducting the MS data analysis. This work was supported by grants from the Japan Society for the Promotion of Science (JP19K16073 and JP21K11673 to MO, JP19KK0195 and JP19H03176 to HT, and JP19H03416 to TO), the Foundation for Promotion of Cancer Research in Japan (to MO), the Hori Sciences and Arts Foundation (to MO), the Sasakawa Scientific Research Grant from the Japan Science Society (to MO), the Takeda Science Foundation (to HT and MO), and the Mitsubishi Foundation (to TO).

## Author Contributions

YT: visualization, investigation, data curation, formal analysis, resources, methodology, validation, writing, review, and editing. MO: visualization, investigation, data curation, formal analysis, funding acquisition, resources, methodology, writing, review, and editing. KY: investigation, data curation, formal analysis. HT: funding acquisition, project administration, supervision, validation, writing, review, and editing TO: conceptualization, data curation, formal analysis, funding acquisition, project administration, supervision, writing-original draft preparation, writing - review, and editing.

## Abbreviations

dEGF20: *Drosophila* Notch EGF20
DLL4: delta-like ligand 4
EGF: epidermal growth factor
ER: endoplasmic reticulum
EOGT: EGF domain-specific O-GlcNAc transferase
FUT: fucosyltransferase
Fuc: fucose
Gal: galactose
GlcNAc: *N*-acetylglucosamine
Hex: hexose
NeuAc: *N*-Acetylneuraminic acid
O-GlcNAc: O-linked *N*-acetylglucosamine

## Data Availability

The glycoproteome data of O-GlcNAc glycan in this study were deposited in jPOST (Project ID: JPST001374).

