## Supplemental Figures for "Comprehensive O-GlcNAc glycoproteomics on NOTCH1 EGF repeats implicated unique Lewis X epitopes in mammals"

**Comprehensive O-GlcNAc glycoproteomics on NOTCH1 EGF repeats refined sequons for O-GlcNAcylation and uncovered unique Lewis X epitopes in mammals**

Yohei Tsukamoto, Mitsutaka Ogawa, Kentarou Yogi, Hideyuki Takeuchi, and Tetsuya Okajima

**A list of the material included**

**Table S1.**

**Table S2.**

**Table S3.**

**Table S4.**

**Figure S1.**

**Figure S2.**

**Figure S3.**

**Figure S4.**

| Vector name | Catalog number |
| --- | --- |
| pCMV6/mouse FUT1: MycDDK | MR218449 |
| pCMV6/mouse FUT2: MycDDK | MR219943 |
| pCMV6/mouse FUT4: MycDDK | MR225818 |
| pCMV6/mouse FUT7: MycDDK | MR225870 |
| pCMV6/mouse FUT9: MycDDK | MR216593 |
| pCMV6/mouse FUT10: MycDDK | MR206957 |
| pCMV6/mouse FUT11: MycDDK | MR223546 |
| pCMV6/mouse POFUT1: MycDDK | MR206140 |
| pCMV6/mouse POFUT2: MycDDK | MR218637 |
| pCMV6/mouse L-FRINGE (LFNG): MycDDK | MR224420 |

**Supplemental Fig S1. Expression vectors for mouse fucosyltransferases (FUTs) and L-Fringe used in this study.**

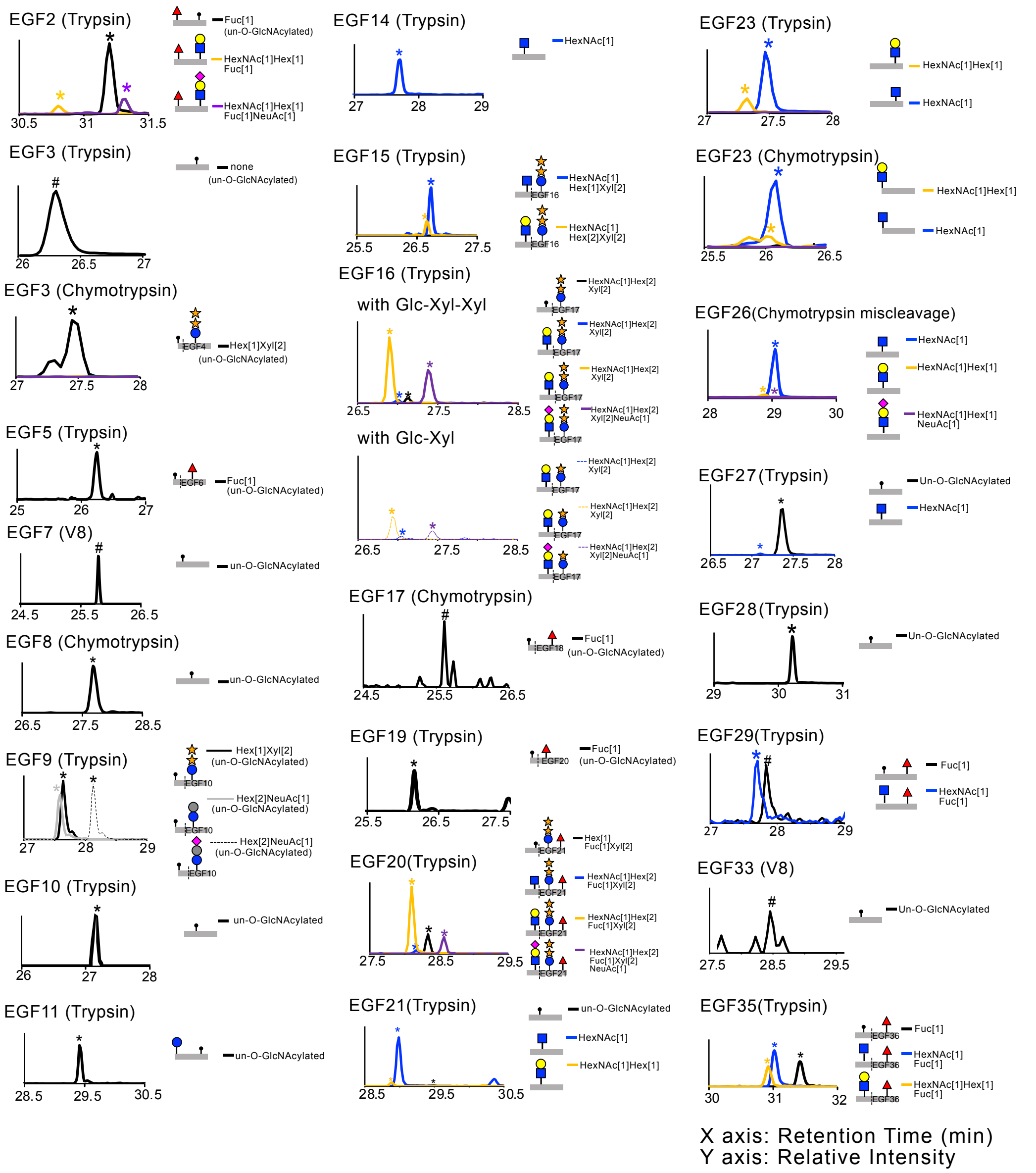


**Supplemental Fig. S2. Semi-quantification of O-GlcNAc glycans on NOTCH1 EGF repeats.** Extracted ion chromatograms (EICs) show the relative ion intensities of protease digests unmodified with O-GlcNAc glycans (*black*) or modified with O-GlcNAc (*blue*), O-GlcNAc-Gal (*orange*), or O-GlcNAc-Gal-NeuAc (*purple*) glycans. The EIC peaks corresponding to the indicated glycoforms determined by the GlycoPAT software or manual inspection are marked by asterisks or hash marks, respectively. It is noted that O-fucose was manually assigned based on the data obtained using different proteases. For example, chymotryptic digestion of full-length NOTCH1 segregates the O-Fuc site at EGF36 from the O-GlcNAc site at EGF35, and the EGF36 O-Fuc site was fully modified with a monosaccharide.

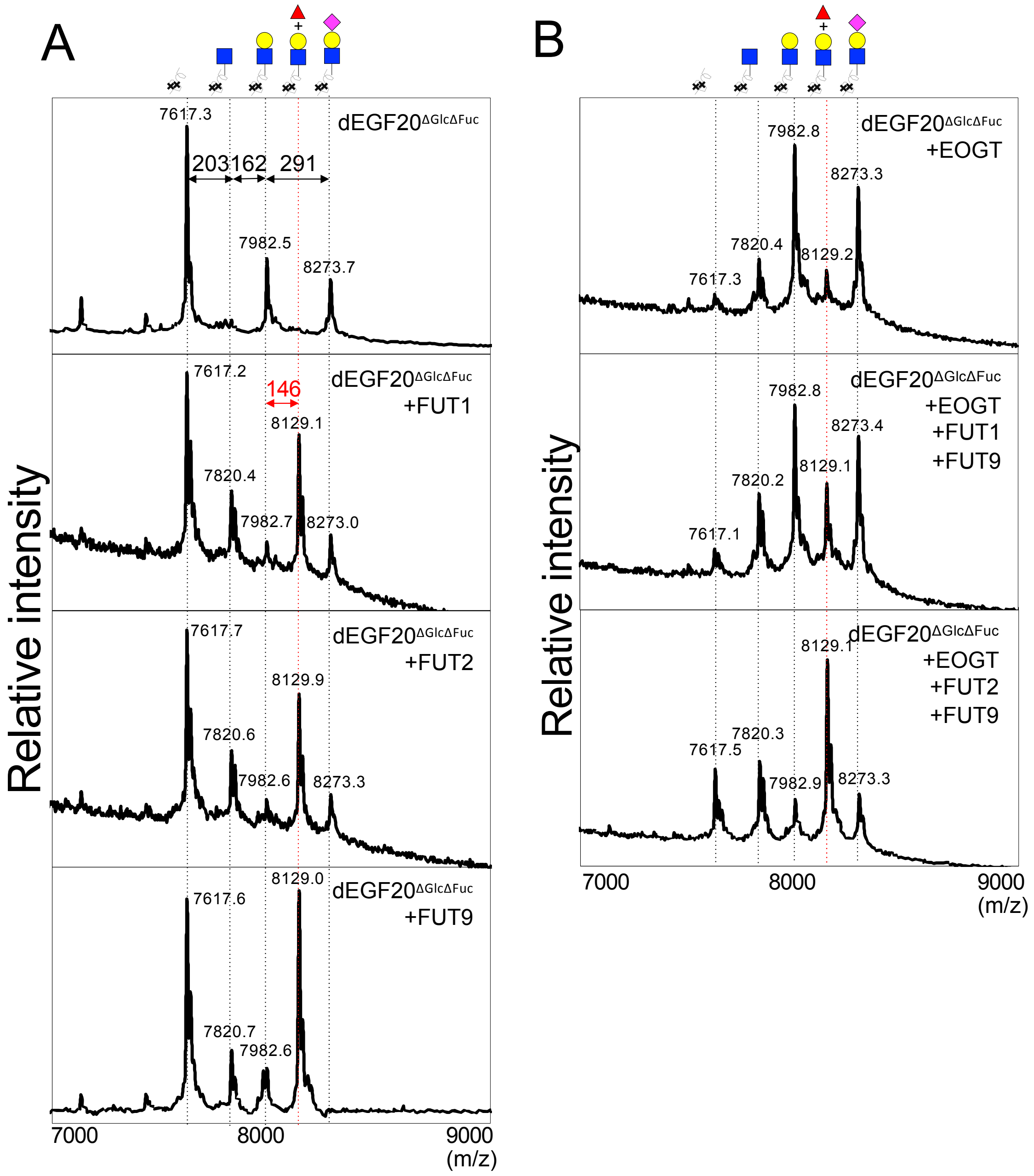


**Supplemental Fig. S3. FUT1, FUT2, and FUT9 create fucose-containing O-GlcNAc glycans on dEGF20. (A)** dEGF20$^{\Delta Glc\Delta Fuc}$-MycHis was transiently expressed in HEK293T cells together with FUT1, FUT2, or FUT9, purified from the culture medium, and analyzed by MALDI-TOF-MS. Also see Figure 2. **(B)** dEGF20$^{\Delta Glc\Delta Fuc}$-MycHis was transiently expressed in HEK293T cells together with EOGT alone, EOGT + FUT1 + FUT9, or EOGT + FUT2 + FUT9 and subjected to MALDI-TOF-MS analysis, as shown above.

**>CXXGYTGXXC**
Human: 48 hit [FAT4, LRP2, HSPG2, TNXB, LAMA1, LAMA2, LAMA3, LAMA5, SVEP1, EYS, CELSR2, FBN3, STAB1, NOTCH1, NOTCH2 NOTCH3, NOTCH4, ACAN, TNC, LAMB2, LAMB4, LAMC3, SLIT2, SNED1, CRB1, MEGF10, MEGF11, PEAR1, MMRN2, DNER, DLL1, DLL4, PAMR1, F12, EGFL6, NTNG2, CLEC18B, CLEC18A, CLEC18C, TMEFF1, TMEFF2, EREG, EPGN]
Mouse: 42 hit [Fat4, Lama1, Lama2, Lama3, Lama5, Hspg2, Svep2, Celsr2, Stab1, Notch1, Notch2, Notch3, Notch4, Tnc, Lamb2, Lamc3, Slit1, Crb1, Crb2, Sned1, Megf9, Megf10, Megf11, Pear1, Nrg2, Dner, Dll1, Dll4, PAMR1, F12, Ntng2, Egfl6, Clec18a, Tmeff1, Tmeff2,Ereg, Epgn]

**>CXXGYSGXXC**
Human: 21 hit [HSPG2, TNXB, RELN, VCAN, LAMA1, LAMA2, NOTCH1, NOTCH2, NOTCH3, CRB1, CRB2, JAG1, JAG2, MMRN1, DLL1, MAC13, DLK1]
Mouse: 16 hit [Lrp2, Hspg2, Svep1, Reln, Vcan, Lama1, Lama2, Notch4, Slit1, Slit2, Slit3, Crb1, Jag1, Egf, Mac13, Mfge8]

**>CXXGFTGXXC**
Human: 38 hit [FAT4, LRP1B, LRP1, HSPG2, TNXB, SVEP1, CELSR1, CELSR2, CELSR3, FBN1, FBN2, NOTCH1, NOTCH2 NOTCH3, NOTCH4, TNC, SNED1, CRB1, TNN, JAG1, JAG2, MEGF10, MEGF11, TIE1, PEAR1, NELL1, NELL2, EMILIN3, DNER, DLL4, VASN, NRG1, COL26A1, NOTCH2NLB, NOTCH2NLC]
Mouse: 34 hit [Fat4, Lrp1b, Lrp1, Svep1, Celsr1, Celsr2, Celsr3, Fbn1, Fbn2, Notch1, Notch2, Notch3, Notch4, Tnc, Megf6, Mfge8, Megf10, Ann, Slit2, Crb1, Sned1, Jag1, Jag2, Lamb3, Tie1, Pear1, Nell1, Nell2, Dner, Dll4, Col26a1]

**>CXXGFSGXXC**
Human: 30 hit [HSPG2, TNXB, SVEP1, RELN, MUC3A, MUC3B, EYS, KALRN, ZAN, STAB1, NOTCH1, NOTCH2, NOTCH3 NOTCH4, STAB2, AGRN, SLIT1, SLIT3, SNED1, JAG1, JAG2, MEGF11, DLL1, DLL4, SELE, DLK1]
Mouse: 18 hit [Hspg2, Svep1, Reln, Notch1, Notch2, Notch3, Agrn, Slit1, Slit3, Sned1, Crb2, Jag1, Megf10, Megf11, Dll1, Dll4, Dlk1]

**>CXXGTTGXXC**
Human: 6 hit [EYS, LAMA2, NOTCH1, NOTCH3, LAMC1, NTNG1]
Mouse: 5 hit [Notch1, Notch3, Lama5, Lama2, Ntng1]

**>CXXGSTGXXC**
Human: 0 hit
Mouse: 0 hit

**>CXXSYTGXXC**
Human: 0 hit
Mouse: 1 hit [Megf6]

**>CXXSYSGXXC**
Human: 1 hit [LRP1B]
Mouse: 1 hit [Lrp1b]

**>CXXSFTGXXC**
Human: 1 hit [NOTCH3]
Mouse: 1 hit [Notch1]

**>CXXSFSGXXC**
Human: 0 hit
Mouse: 0 hit

**>CXXSTTGXXC**
Human: 0 hit
Mouse: 0 hit

**>CXXSTSGXXC**
Human: 0 hit
Mouse: 0 hit

**>CXXPYTGXXC**
Human: 5 hit [FAT2, NOTCH1, NOTCH2, TNR, HABP2]
Mouse: 6 hit [Notch1, Notch2, Fat2, Habp2, Crb1, Tnr]

**>CXXPYSGXXC**
Human: 0 hit
Mouse: 0 hit

**>CXXPFTGXXC**
Human: 3 hit [EYS, NOTCH1, MMRN1]
Mouse: 0 hit

**>CXXPFSGXXC**
Human: 4 hit [EYS, NOTCH2, FBXL18, HABP2]
Mouse: 1 hit [Habp2]

**>CXXPTTGXXC**
Human: 0 hit
Mouse: 0 hit

**>CXXPTSGXXC**
Human: 0 hit
Mouse: 0 hit

**Supplemental Fig. S4. List of glycoproteins predicted to be O-GlcNAcylated by EOGT.** The refined sequons for O-GlcNAcylation defined as $C^5$-X-X-G/S/P-Y/F/T-T/S-G-X-X-$C^6$ predict approximately 100 proteins that are modified with O-GlcNAc by EOGT.

**Supplemental Table S1. Orbitrap Fusion Tribrid MS acquisition parameters.** AGC, automatic gain control; NCE, normalized collision energy.

**Supplemental Table S2. List of O-GlcNAcylated or un-O-GlcNAcylated peptides detected by the GlycoPAT software.** Where indicated, fucose was assigned on the O-Fuc site as a fixed monosaccharide modification, based on data from other glycopeptides containing O-Fuc sites but lacking O-GlcNAc sites. Scan, the scan number of the MS/MS spectrum; Expt, experimental mass of the precursor ion; Mono, monoisotopic mass of the candidate glycopeptide; Charge, precursor ion charge state; Percent Ion Match, % of ions matched among all theoretical fragments; Top10, number of Top 10 peak matched; Ensemble Score, a weighted average parameter of four different statistical metrics including Xcorr, % ion match, *p* value, and Top10 peaks.

**Supplemental Table S3. List of annotated MS/MS spectra for O-GlcNAcylated or un-O-GlcNAcylated peptides detected by the GlycoPAT software and manual inspection.** In an annotated spectrum generated by the GlycoPAT software, *green* denotes peaks matched by fragment ions of the candidate peptide; *red* indicates peaks that could not be matched. Blue color shows manually annotated glycan-derived oxonium ions.

**Supplemental Table S4. List of O-GlcNAcylated or un-O-GlcNAcylated peptides detected by manual inspection.**
